# Artificial symbiont replacement in a vertically-transmitted plant-bacterium association provides insights into the basis for specificity

**DOI:** 10.1101/2025.03.10.642324

**Authors:** Léa Ninzatti, Thibault G. Sana, Tessa Acar, Sandra Moreau, Marie-Françoise Jardinaud, Guillaume Marti, Olivier Coen, Aurélien Carlier

## Abstract

Some plants engage in permanent, vertically transmitted symbioses with bacteria. Often, these bacteria are hosted extracellularly in specific structures on the leaves, where they produce specialized bioactive metabolites that benefit their host. These associations are highly specific, with one plant species associating with a single bacterial species, but little is known about how these symbioses originate and how specificity is maintained. In this study, we show that the symbiotic association between a wild yam and a bacterium can be manipulated experimentally, and that bacteria-free plants are open to colonization by environmental bacteria. Through metabolic profiling, we show that the endophytic niche is rich in organic acids and intermediates of the TCA cycle. Environmental bacteria capable of utilizing these acids, such as the soil bacterium *Pseudomonas putida*, readily colonize aposymbiotic plants. However, successful colonization is contingent upon the absence of the vertically-transmitted symbiont and an active type VI secretion system. Unexpectedly for a vertically-transmitted symbiosis, these findings suggest that microbe-microbe interactions, including antagonism, may play a crucial role in maintaining the specificity of an association. However, low transmission rates of synthetic symbionts provide evidence that transmission barriers or bottlenecks may still occur, further enforcing partner fidelity. Together, these results highlight the complexity of mechanisms underlying mutualistic associations, and provide insights into the evolution of bacterial leaf symbiosis.

## Introduction

Mutualistic interactions are generally costly to maintain, and a degree of specificity is often important for the maintenance of a beneficial holobiont. Specificity between symbiotic partners may be driven by a combination of three factors: the mode of acquisition of the symbiont, environmental selection (e.g. via restrictive nutrients or physico-chemical properties within the niche), and mechanisms underlying recognition between the host and its symbiont [1]. For example, in well-studied model systems such as the legume-*Rhizobium* and the squid-*Vibrio* symbioses, partner specificity is achieved by a combination of an aposymbiotic phase, during which the host exists without the symbiotic partner, and selective barriers allowing the establishment of a cooperative symbiont [2, 3]. Microbe-microbe interactions may also play a key role in the establishment of a symbiosis, where a host may foster interference between a symbiont and non-productive competitors [4].

Interference competition as a mechanism underlying partner choice has been experimentally demonstrated in systems in which hosts acquire bacteria from the environment at the juvenile stage such as the attine ants/*Streptomyces* and the *Riptortus pedestris/Burkholderia* symbioses [5, 6]. Finally, some symbionts are also transmitted vertically from mother to offspring. Vertical transmission links the evolutionary fates of the host and symbionts, which promotes the evolution of mutualistic interactions through partner-fidelity feedback and a high degree of symbiont relatedness within a host [7].

Moreover, vertical transmission usually occurs at a single stage of the lifecycle (e.g. by infecting the germline), thus sheltering symbionts from competition [8]. Strictly vertically-transmitted symbionts do not need to be selected from an environmental population, and sophisticated mechanisms to enforce specificity are thought not to be necessary.

Hereditary symbiosis is common in animals, especially insects, but much less described in plants. However, some plants engage in permanent, strictly specific associations with bacteria within their leaves, a phenomenon known as leaf symbiosis. Leaf symbiosis has been described in about 500 plant species, mostly in the Primulaceae and Rubiaceae families [9, 10]. The bacteria, mainly belonging to the *Burkholderia sensu lato*, are present in leaf nodules that can take various shapes [11]. Several lines of evidence indicate that the leaf symbionts of some species have a defensive role. For example, *Candidatus* Caballeronia kirkii (*Ca*. C. kirkii); the symbiont of *Psychotria kirkii* (Rubiaceae), is linked to the accumulation of bioactive cyclitols in the plant. Of these, kirkamide displays cytotoxic and insecticidal properties [12], and streptol glucoside is a potent herbicide [13]. The bacterial symbionts are passed on to the next host generation via the seed [10, 14], but how the symbiotic bacteria reliably colonize seedlings in the presence of a complex soil and seed microbiome remains unknown.

The yam *Dioscorea sansibarensis* forms a permanent association with the bacteria *Orrella dioscoreae*, and offers an experimentally tractable system of heritable leaf symbiosis [15, 16]. The interaction most obviously takes place in a prominent gland at the tip of the leaf. We recently demonstrated that *O. dioscoreae* is vertically transmitted through vegetative propagules, called bulbils, and therefore present throughout the entire plant lifecycle [17].

Co-phylogenetic analyses indicate that the transmission route is not entirely closed, with horizontal transmission or host-switching also occurring in the wild [18]. However, the *D. sansibarensis-O. dioscoreae* symbiosis is ubiquitous in nature, implying the existence of mechanisms to enforce specificity despite evidence of a permissive transmission route [18]. In the laboratory, the plant and the bacteria can be grown separately, suggesting that this symbiosis is not obligate despite its specificity in nature. Remarkably, aposymbiotic plants develop normally *in vitro*, with fully formed leaf glands devoid of bacteria [19].

We demonstrated in a previous study that infection of aposymbiotic plants relies on colonization of the shoot apical bud, which contains the meristematic tissue at the origin of all above-ground organs development throughout the life of the plant [17]. In this regard, plants face a drastically different situation from animals regarding vertical transmission of symbionts. Because the meristem links somatic and reproductive organs, plants cannot easily segregate a symbiont population between a germ line and a somatic population.

Moreover, our previous studies showed that the apical bud remains physically open to the environment, creating opportunities for other microorganisms to establish in the plant at virtually any point during the growing phase [19]. What mechanisms evolved to enforce partner specificity in this context is unknown.

The presence of *O. dioscoreae*within the plant may prevent strains with similar niche requirements from establishing in the plants via priority effects, for instance by pre-empting access to favored carbon or nitrogen sources. Priority effects may also be paired with other mechanisms such as interference competition, to ensure higher specificity. Type VI secretion systems (T6SSs) are known to have anti-microbial functions through delivery of effectors directly into targeted cells in a contact-dependent manner, and play a key role in microbe-microbe interactions [20, 21]. Indeed, the genome of *O. dioscoreae* harbors two T6SS gene clusters that are highly expressed in the leaf gland and conserved in all genomes of *O. dioscoreae* sequenced so far [18, 22]. Both gene clusters code for the 13 core proteins: TssABCEFGJKLM, VgrG (Valine-glycine repeat G protein), Hcp (Haemolysin co-regulated protein) and PAAR (Proline-Alanine-Alanine-Arginine domaine containing protein), as well as for accessory proteins which usually play a role in regulation and modulation of the system assembling [23].

T6SSs are composed of 4 main parts, the integral membrane complex (TssJL and M), the baseplate (TssAEFG and K), the spike (Hcp, VrgG and PAAR) and the contractile sheath (TssBC). The spike is constituted of Hcp proteins polymerized into a stack of hexameric ring, topped by the VgrG trimer and the PAAR protein [24–26]. The contraction of the sheath ejects the spike into target cell, delivering the effector proteins (Zoued et al. 2016; Leiman et al. 2009; Basler et al. 2012; Wang, Brodmann, et Basler 2019). The membrane complex is stable and may be used for multiple injections after reassembly. In the contracted conformation the ClpV AAA+ ATPase is recruited to depolymerize the TssBC subunits, allowing recycling of the system [31, 32]. Interfering with the ClpV ATPase usually results in specific inactivation of the cognate T6SS, preventing recycling of the system after effector delivery.

The role of T6SS-mediated competition in niche monopolization has been demonstrated in various Gram-negative bacteria. Notably, T6SSs were first described in the pathogenic bacterium *Vibrio cholerae* as essential for both colonization and virulence [20, 33]. Specifically, T6SS-mediated competition is crucial for *V. cholerae* to outcompete resident microbiota in the human gut during disease [34].

The aim of this study was to acquire a better understanding of the mechanisms underlying specificity in the *D. sansibarensis-O. dioscoreae* symbiosis. We show that despite a ubiquitous and highly specific association, leaf glands of aposymbiotic *D. sansibarensis* are open to benign colonization by unrelated bacterial species with similar niche requirements. Co-inoculation assays on aposymbiotic seedlings show that *O. dioscoreae* is highly competitive *in planta*, and that this competitiveness is dependent upon a functional T6SS.

## Material and methods

### Plant culture and propagation

*Dioscorea sansibarensis* Pax plants were obtained from the greenhouse of the Botanical Garden at the University of Ghent (LM-UGent) in Ghent, Belgium. Chemicals and reagents were purchased from Merck unless otherwise indicated. Plants used throughout in experiments were maintained in the greenhouse of the Laboratory of Plants Microbes and Environment Interactions (LIPME) in Castanet-Tolosan, France. Unless otherwise indicated, plants were grown in climate-controlled chambers at 28°C, 70% humidity and a light cycle of 16h light (210 μmol/m^2^/s), 8h dark.

### Production of aposymbiotic plants

Aposymbiotic plants were produced from node cutting treated with antibiotics as described in [17]. Briefly, nodes were dissected from growth chamber-grown plants, and surface sterilized for 8 hrs in a solution of 3X Murashige Skoog medium (MS, Sigma) supplemented with 5% of Plant Preservative Mixture (PPM, Plant Cell Technology, USA). Explants were then aseptically transferred to sterile 6-well plates containing MS medium supplemented with sucrose 2% (Merk), myo-inositol 555 µM (Sigma), glycine 26.6 µM (Sigma), cysteine 16.5 µM (Sigma), nicotinic acid 4.06 µM (Sigma), pyridoxine 2,96 µM (Sigma), thiamine 1,88 µM (Sigma), PPM (0.2% w/v) and the antibiotics carbenicillin and cefotaxim (200 µg/mL each). Explants were incubated in a growth chamber at 28°C under 16h/8h day/night cycle. Medium was replaced after 10 days. After 3 weeks of incubation, the explants were aseptically transferred to sterile Magenta boxes (model GA7, Merck) containing MS medium as above without the carbenicillin and cefotaxim. Effectiveness of the treatment was tested for each explant by collecting the first 2 leaves, milling in 100 µl of 0.4% w/v NaCl using a Restch MM400 bead mill (1 minute, 30Hz), and spreading the macerate on TSA medium (Millipore). The absence of visible microbial growth after 2 days of incubation at 28°C was taken as evidence of the aposymbiotic status of the plants.

### Isolation and identification of leaf gland bacteria

Aposymbiotic plants were transferred from closed, sterile containers to open pots in greenhouse (25°C, 16h light/8h dark, 60% humidity). Leaf acumens were dissected and surface sterilized (5 min. in 70% ethanol, sterile distilled water wash, 5 min. in 1,6% Bleach, 3 washes in sterile NaCl 0.4%). Samples were homogenized as described above, and plated on TSA medium. Bacterial colonies were isolated on TSA. Identification was done by 16S rRNA gene sequencing using Sanger sequencing and PCR primers 27f and 1492r [35].

### Inoculations of *D. sansibarensis* aposymbiotic plant with bacteria

Overnight bacterial cultures were grown in the appropriate medium and harvested by centrifugation at 7000 g for 5 min. Cell pellets were washed twice in sterile 0.4% NaCl. Bacterial suspensions were normalized to OD_600nm_ = 0.2 for each culture. Prior to inoculation, plants were moved to sterile Microbox containers (SacO2, Belgium) containing 50 mL of MS medium as above without the carbenicillin and cefotaxim. Plants were inoculated by depositing 2 µL of the bacterial suspension directly onto the apical bud.

Inoculated plants were grown at 28°C with a 16h light/8h dark cycle for 28 days. When applicable, a second inoculation was done a week later as described above. Colonization was evaluated by collecting leaf glands, and spreading serial dilutions of the milled and weighted acumens onto appropriate media as described above.

### Detection of bacteria in the second generation of inoculated plants

Previously inoculated plants were transferred from closed containers to open pots in growth chambers (25°C, 16h light/8h dark, 70% humidity) until the end of the growing season.

Bulbils were harvested and stored in a dry place until spontaneous sprouting. Sprouting bulbils were laid on a moist sand substrate and transferred to a growth chamber with high hygrometry (25°C, 16h light/8h dark, 90% hygrometry). Plantlets with at least 12 leaves they were transferred to lower hygrometry chamber (25°C, 16h light/8h dark, 65-70% hygrometry). Two acumens per plant were tested for the presence of the bacteria. Leaf acumens were dissected and surface-sterilized (5min 70% ethanol, 3 min 1,4% Bleach) and the presence and identity of bacteria was tested as describe above.

### Observation of bacteria in leaf glands by fluorescent microscopy

For the observation of fluorescent bacteria, leaf glands were dissected with sterile scissors 8 weeks after inoculation and fixed in 0.3% paraformaldehyde in 0.1 M pH7 potassium phosphate buffer for two hours at room temperature under vacuum. Samples were rinsed twice with potassium phosphate buffer (0.1 M, pH7). One hundred µm-thick Sections were prepared using a vibrating microtome (Leica VT1000S) after embedding in 5% low melting agarose (Nusieve). Sections were observed using an epifluorescence microscope (Zeiss Axioplan2). Images were processed using the ImageJ software version 1.54.

Observation of non-fluorescent bacteria was done by collecting leaf glands 5 weeks post inoculation. Samples were prepared by sectioning with a razor blade, followed by staining with Syto9 (Thermo Fischer) and visualization using confocal microscope (Leica SP7). Image were processed using Leica LasX software.

### Bacterial strains, plasmids and growth conditions

Bacterial strains and plasmids are listed in Table S1. Routine culture of *O. dioscoreae* strains was done at 28°C in Tryptic Soy Broth (TSB, Sigma) supplemented with 30 µg/ml of nalidixic acid, 50 µg/ml of kanamycin or 20 µg/ml of gentamicin where appropriate.

*Stenotrophomonas sp.*, *Rhizobium sp.* and *Sphingomonas sp.* strains were grown in TSB medium at 28°C unless otherwise specified. *P. putida* and *E. coli* strains were grown in LB medium with appropriate antibiotics. To selectively grow *O. dioscoreae* R-71412 and derivatives, AB minimal medium [36] was supplemented with 10 mM trisodium citrate (Sigma), nalidixic acid (30 µg/ml) and gentamicin (20 µg/ml). Pseudomonas Isolation Agar (MilliPore) supplemented with 2% glycerol was used to grow *P. putida* selectively.

### Bacterial genetics

T6SS deletion mutants were produced in an R-71412 background by homologous recombination as described in Sana et al. 2024, using a protocol inspired by [38]. Briefly, the flanking regions of the gene to delete were amplified by PCR using primers listed in Table S2. The pSNW2 plasmid DNA was digested by XmaI (New England Biolabs) and gene fragments were assembled using the Pro Ligation-free cloning kit (Applied Biological Materials, Richmond, BC, Canada) and introduced into *E. coli* DB3.1 *λpir* using standard heat-shock protocols. All constructs were verified by whole plasmid sequencing using ONT Nanopore sequencing [39]. *O. dioscoreae* R-71412 was electroporated purified plasmid DNA as previously, and kanamycin-resistant clones were selected [17]. Conditionally-replicating plasmid pQURE6 harboring a I-SceI nuclease gene was introduced into merodiploid clones by triparental mating. Colonies with gene replacement events were screened for loss of kanamycin resistance and by PCR. The double mutant was produced by deletion of ODI_R0808 in the previously obtained ODI_R3997 deletion mutant. The genome sequences of all mutant strains were obtained by Oxford Nanopore sequencing using R10.4 chemistry on an ONT P2 solo instrument.

For genetic complementation, gene fragment were amplified by PCR from *O. dioscoreae* R-71412 genomic DNA using primeSTAR Max DNA polymerase (Takara Bio) with primers listed in Table S2. Purified PCR product were cloned into plasmid pSEVA2313 by restriction and ligation. *E. coli* Top10 was transformed by electroporation, plasmids were isolated from selected clones and validated by sequencing as above. Purified plasmid DNA was introduced into *O. dioscoreae* strains as above. Transformants were selected on TSA medium supplemented with kanamycin (50 mg/L).

### Bacterial phenotyping

Phenotyping Microarray (PM) plates were purchased from Biolog (USA). Bacteria were grown on solid media containing appropriate antibiotics and resuspended into sterile distilled water as per the manufacturer’s recommendation. Trisodium citrate was added to wells of the PM03 plate to 0.5% w/v final concentration. NADH oxidation was measured every 15 minutes for 120 hours using an OmniLog instrument. Data were analyzed with the opm R package [40]. For quantitative comparison of *O. dioscoreae* R-71412 and *P. putida* KT2440*::gfp* metabolism of select carbon sources, custom Biolog plates were prepared with AB medium supplemented with either trisodium citrate, L-malate, sodium succinate, fumarate or sodium pyruvate to 10 mM final concentration. Data were analyzed on R using the Growthcurver package [41]. Optimal pH range for growth was tested in LB medium supplemented with 10 mM of trisodium citrate adjusted to pH = 5, 6, 7 or 8 and buffered with 100 mM of MES, MOPS or EPPS buffer as appropriate; non-adjusted (pH 7) LB- trisodium citrate medium was used as control. Cultures were inoculated at 1:100. Bacterial growth was monitored by OD_600_ readings every 30 minutes for 48h, using BMG FLUOstar Omega microplate reader. Growth rate was calculated on R using the Growthcurver package [41].

### *In vitro* bacterial competition

LB media was inoculated with a suspension of *O. dioscoreae* mCherry-tagged strain R-71417 and/or with *P. putida* KT2440::*gfp* strain in sterile distilled water normalized at OD_600nm_ = 0.02. The R-71417 was co-inoculated at the same time as the KT2440::*gfp*, or prior (2h, 4h or 6h) to the inoculation of the KT2440::*gfp*. Total bacterial growth was measured at OD600. Relative growth of *O. dioscoreae* R-71417 and *P. putida* KT2440::*gfp* was estimated by fluorescence measurement, respectively with mCherry (544nm/590nm) and GFP (485nm/520nm) filters. Experiments were done using BMG FLUOstar Omega microplate reader for 24 hours with a measure taken every hour. Growth rate was calculated using the Growthcurver R package. Fluorescence intensities were normalized relative to the mean fluorescence of the monoculture in each replicate.

### *In vitro* contact-dependent competition assay

Fluorescence-dependent killing assays of *O. dioscoreae* R-71412, T6SS mutants and complemented strains, as well as fluorescent *P. putida* KT2440::*gfp* were grown overnight in TSB with appropriate antibiotics at 28°C. Cells were washed in a sterile solution of NaCl 0.4% w/v and density was normalized to OD_600nm_ = 0.5. Then 10 µl of each sample were spotted onto a 96-well plate (Nunc FluoroNunc) filled with 180 µl of solid growth medium (TSA).

Fluorescence was measured for 24h at 28°C, in a BMG FLUOstar Omega microplate reader (set for spiral top reading, 30 reading points, adjusted with 90% of a 900 arbitrary fluorescence unit gain, filters: excitation 485-12nm and emission 520-20nm). For cfu-counting, cultures of *O. dioscoreae* and *P. putida* strains were prepared as above and concentrated in sterile 0.4% NaCl solution to OD_600_ =50 for *O. dioscoreae* strains and OD_600_ =10 for *P. putida*. Then 5 µl of each sample were spotted onto pre-warmed TSA. After 4h at 28°C, cells were resuspended in 1 ml of NaCl 0.4% (w/v), serially diluted and plated on PIA and AB media supplemented with 10 mM citrate with appropriate antibiotics to estimate the number of *P. putida* and *O. dioscoreae* colony forming units (cfu), respectively.

### Ultra High Performance Liquid Chromatography High Resolution Mass Spectrometry analysis

Acumens from aposymbiotic plants re-inoculated with *O. dioscoreae* R-71412 or with a NaCl 0.4% solution were dissected, and immediately frozen in liquid nitrogen. Samples were milled using a Restch MM400 bead mill (30sec, 30Hz), and stored at -80°C until extraction. Samples were extracted with a mixture of methanol:water 80:20, with a proportion of 1ml of solvent per 100 mg of sample. Ultra-high-performance liquid chromatography-high-resolution MS (UHPLC-HRMS) analyses were performed on a Q Exactive Plus quadrupole (Orbitrap) mass spectrometer, equipped with a heated electrospray probe (HESI II) coupled to a U-HPLC Ultimate 3000 RSLC system (Thermo Fisher Scientific, Hemel Hempstead, United Kigdom). Separation was done on a Luna Omega Polar C18 column (150 mm × 2.1 mm i.d., 1.6 μm, Phenomenex, Sartrouville, France) equipped with a guard column. The mobile phase A (MPA) was water with 0.05% formic acid (FA), and the mobile phase B (MPB) was acetonitrile with 0.05% FA. The solvent gradient was 0 min, 100% MPA; 1 min, 100% MPA; 22 min, 100% MPB; 25 min, 100% MPB; 25.5 min, 100% MPA; and 28 min, 100% MPA.

The flow rate was 0.3 ml/min, the column temperature was set to 40°C, the autosampler temperature was set to 5°C, and the injection volume was fixed to 5 μl. Mass detection was performed in positive ionization (PI) mode at resolution 35,000 power [full width at half- maximum (fwhm) at 400 m/z] for MS1 and 17,500 for MS2 with an automatic gain control (AGC) target of 1 × 106 for full scan MS1 and 1 × 105 for MS2. Ionization spray voltages were set to 3.5 kV, and the capillary temperature was kept at 2561C. The mass scanning range was m/z 100-1500. Each full MS scan was followed by data-dependent acquisition of MS/MS spectra for the six most intense ions using stepped normalized collision energy of 20, 40, and 60 eV.

The raw data were processed with MSdial version 4.70 for mass signal extraction between 100 and 1,500 Da [42]. MS1 and MS2 tolerance were set to 0.01 and 0.025 Da in the centroid mode. The optimized detection threshold was set to 5 × 10^5^ concerning MS1 and 10 for MS2. Peaks were aligned on a QC reference file with a retention time tolerance of 0.15 minutes and a mass tolerance of 0.015 Da. Peak annotation was performed with an in- house database built on an MS-FINDER model [43]. MS-DIAL data were then cleaned with the MS-CleanR workflow by selecting all filters with a minimum blank ratio set to 0.8, a maximum relative standard deviation (RSD) set to 30, and a relative mass defect (RMD) ranging from 50 to 3.000. The maximum mass difference for feature relationships detection was set to 0.005 Da and the maximum RT difference to 0.025 min. Pearson correlation links were considered with correlation ≥ 0.8 and statistically significant with α = 0.05. Two peaks were kept in each cluster, viz., the most intense, and the most connected. The kept features (m/z × RT pairs) were annotated with MS-FINDER version 3.52. The MS1 and MS2 tolerances were, respectively, set to 10 and 20 ppm. Formula finders were only processed with C, H, O, N, and S atoms. Databases (DBs) based on Dioscorea (genus), Alcaligenaceae (family) were constituted with the dictionary of natural products (DNP, CRC press, DNP on DVD v. 28.2). The internal generic DBs from MS-FINDER used were KNApSAcK, PlantCyc, NANPDB, UNPD, COCONUT, and CheBI. Statistical analyses were done using the R software and standard packages.

## Results

### Aposymbiotic *D. sansibarensis* are open to colonization by bacteria other than *O. dioscoreae*

We reported previously that aposymbiotic *D. sansibarensis* plants were amenable to colonization by exogenously applied *O. dioscoreae*, suggesting that aposymbiotic plants remain receptive to bacteria after germination. To test whether this applied to bacteria other than *O. dioscoreae*, we produced bacteria-free plants by antibiotic treatment of explants. After approximately 2 months in sterile cultures, plants were moved to open pots in the greenhouse for the rest of the growing season. The leaf glands of several of those aposymbiotic plants became spontaneously colonized with bacteria unrelated taxonomically to *O. dioscoreae* (Table 1). To test if these strains reliably colonized *D. sansibarensis* leaf glands, we selected 3 isolates belonging to the genera *Stenotrophomonas sp.* (R-67087)*, Rhizobium sp.* (R-71694) and *Sphingomonas sp.* (R-71695) for further study. Artificial inoculation of aposymbiotic plants with these three strains resulted in reliable colonization of leaf glands, albeit with lower densities than observed for *O. dioscoreae* (Figure 1).

**Figure 1.**
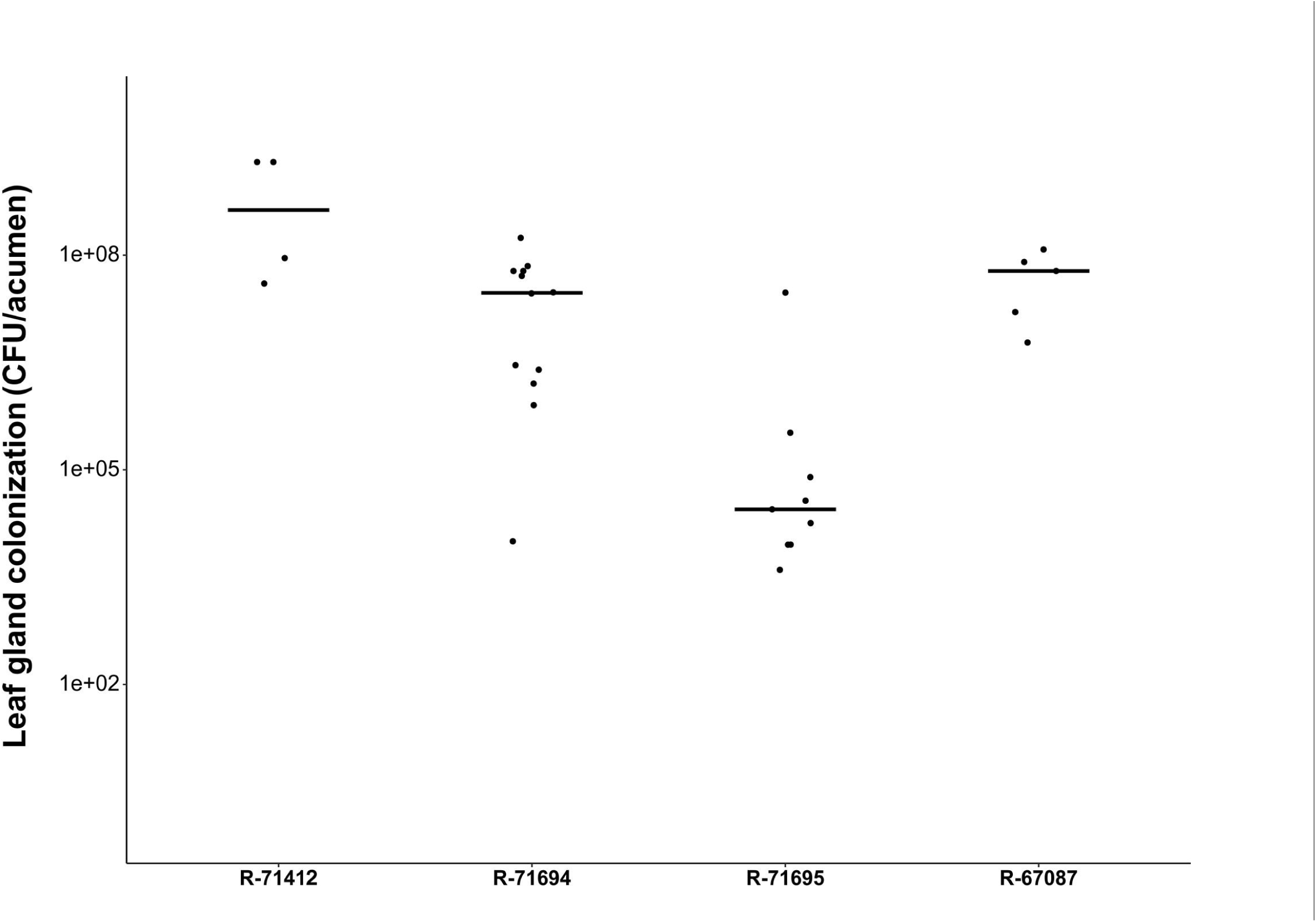
Non-symbiotic bacteria effectively colonize *D. sansibarensis* leaf glands. Aposymbiotic *D. sansibarensis* were inoculated on the apical meristem with *O. dioscoreae* (R-71412), *Rhizobium sp.* (R-71694), *Sphingomonas sp.* (R-71695) and *Stenotrophomnas sp.*(R-67087). Newly emerged acumens were macerated and serial dilutions were plated out to yield an estimate of cfu/leaf acumen.

**Table 1.** Identification of isolates from glands of aposymbiotic *D. sansibarensis*.

| **Number of plants**<br>**colonized** | **Species (best blast hit)** | **Best Hit** | **Similarity (%)** |
| --- | --- | --- | --- |
| 5 | *Rhizobium radiobacter* | ATCC 19358 | 99.68 |
| 1 | *Cupriavidus plantarum* | ASC-64 | 99.44 |
| 1 | *Variovorax* sp. | PMC12 | 100 |
| 1 | *Pantoea* sp. | ND03 | 99.86 |
| 1 | *Pseudomonas atacamensis* | M7D1 | 99.79 |
| 1 | *Enterobacter soli* | ATCC BAA-2102 | 99.71 |
| 1 | *Agrobacterium* sp. | K599 | 100 |
| 2 | *Stenotrophomonas indicatrix* | WS40 | 100 |
| 1 | *Pseudomonas fortuita* | GMI12077 | 100 |
| 2 | *Pseudomonas lactis* | DSM 29167 | 100 |
| 1 | *Pseudomonas citronellolis* | NBRC 103043 | 100 |
| 1 | *Pseudomonas neuropathica* | P155 | 99.92 |
| 1 | *Pseudomonas paralactis* | DSM 29164 | 99.51 |
| 1 | *Cupriavidus campinensis* | WS2 | 99.71 |

**Table 2.** Detection of metabolites supporting the growth of *O. dioscoreae* in symbiotic and aposymbiotic leaf glands of *D. sansibarensis*. Average peak intensity of metabolites detected in the symbiotic and aposymbiotic leaf glands of *D. sansibarensis*. Statistical significance of the fold change was calculated by Student’s T-test. Standard deviations are indicated for each average intensity value. ID” of Table S3 and Table S4.

| Metabolite | Feature ID* | Peak Intensity |  | Fold change | Student's T-test p-value |
| --- | --- | --- | --- | --- | --- |
|  |  | Aposymbiotic | Symbiotic |  |  |
| Citrate | 162_C18neg | 7310.65 ± 3646.79 | 4022.14 ± 2342.12 | 1.82 | 0.087 |
| Gluconate | 175_C18neg | 330.21 ± 124.02 | 233.70 ± 140.22 | 1.42 | 0.133 |
| Aspartic acid | 63_C18neg | 487.17 ± 169.64 | 299.37 ± 123.06 | 1.63 | 0.056 |
| L-pyroglutamic acid | 51_C18neg | -34.03 ± 3.62 | -39.65 ± 9.57 | 0.86 | 0.091 |
| Malate | 66_C18neg | 3520.54 ± 909.21 | 1760.72 ± 534.42 | 2.00 | 0.012 |
| L-Glutamine | 83_C18neg | 145.21 ± 71.42 | 249.63 ± 76.65 | 0.58 | 0.041 |
| L-glutamic acid | 84_C18neg | 391.37 ± 54.89 | 427.43 ± 219.53 | 0.92 | 0.355 |
| Galactonic acid gamma-lactone | 323_C18pos | -35.94 ± 0.91 | -3.71 ± 19.26 | 9.69 | 0.002 |
| L-Serine | 4_C18neg | 875.24 ± 370.06 | 206.85 ± 140.09 | 4.23 | 0.016 |

Histological staining of the leaf glands demonstrated that all three strains occupied the lumen of the leaf gland, similar to *Orrella dioscoreae* R-71412 (Figure S1).

### Characterization of the metabolic niche of *O. dioscoreae*

The diverse taxonomic range isolates from aposymbiotic leaf glands prompted us to test whether bacteria with similar physiological properties to *O. dioscoreae*might be able to colonize aposymbiotic *D. sansibarensis*. *O. dioscoreae* grows aerobically, requires a range of temperatures for growth of 15-40°C and pH slightly acidic to neutral (Figure S2). We screened a library of compounds to get a comprehensive overview of potential nutrients available for growth by *O. dioscoreae*. Of the 190 carbon sources tested, only 27 were oxidized by cultures of *O. dioscoreae* after 72 hours (Table S1). Notably, oxidation of citrate, malate, succinate and fumarate indicated that assimilation pathways converge towards the tricarboxylic acid cycle (TCA). Hexose sugars were not utilized by *O. dioscoreae*, and gluconate supported growth for some, but not all, strains [16]. We also screened 95 compounds as potential sources of nitrogen (Table S1). In particular, ammonia, urea and the amino acids L-Ala, L-Asn, L-Asp, L-Glu, L-Gln, Gly, L-His, L-Ile, L-Leu, L-Pro, L-Ser, L-Met, D-Ala, D-Asp, D-Glu, D-Ser supported cellular activity as nitrogen sources. Metabolomics analysis of symbiotic vs. aposymbiotic glands confirmed that malate, citrate aspartate and glutamate were abundant in whole leaf glands. Strikingly, malate was significantly enriched by nearly 2-fold in aposymbiotic vs. symbiotic leaf glands (Student t-test *p*-value = 0.012).

Concentrations of citrate were not significantly different, but tended to be somewhat more abundant in extracts of aposymbiotic leaf glands, (1.82x, p-value = 0.087) (Table 3). We were unable to detect unambiguously other intermediates of the TCA in our samples, but the fact that citric acid and malic acid are slightly depleted when bacteria are present within the leaf glands indicated that these compounds may be metabolized by the bacteria within the leaf gland.

**Table 3.** Detection of bacteria in offspring of inoculated *D. sansibarensis*.

| Inoculum | *P. putida* | *P. putida* + *O. dioscoreae* |
| --- | --- | --- |
| Average number of bulbils produced by plant | 5.9 | 4.4 |
| Sprouting efficacy | 9.1% | 15.7% |
| Detection of only *P. putida* in offspring | 0 | 0 |
| Detection of only *O. dioscoreae* in offspring | 0 | 68 (62.3%) |
| Detection of *O. dioscoreae* and *P. putida* | 0 | 1 (0.9%) |
| No bacteria detected | 35 | 40 (36.7%) |
| Total number of tested acumens | 35 | 109 |

### Metabolic niche overlap between *P. putida* and *O. dioscoreae*

We searched the available literature for bacterial strains capable of utilizing all the identified carbon sources of *O. dioscoreae*. *Pseudomonas putida* KT2440 is a well described model organism capable of growth on many organic acids, and is derived from *P. putida* mt-2 originally isolated from soil [44, 45]. The metabolic requirements of strain KT2440 are well documented and comprise those of *O. dioscoreae* [46–50](Table S2). First, we confirmed that a GFP-tagged derivative of *P. putida* KT2440 could utilize all the carbon and nitrogen sources that support the growth of *O. dioscoreae* (Table S1). Next, we measured the respiration of *O. dioscoreae* and *P. putida* KT2440::*gfp* on five substrates: trisodium citrate, L-malate, sodium succinate, fumarate and sodium pyruvate. Cultures of *P. putida* KT2440::*gfp* showed higher respiration rate (µ) and maximum curve height (A) with L-malate, trisodium citrate and sodium succinate than *O. dioscoreae* (Figure 2). We detected no differences between the 2 strains regarding the oxidation of fumarate or sodium pyruvate.

**Figure 2.**
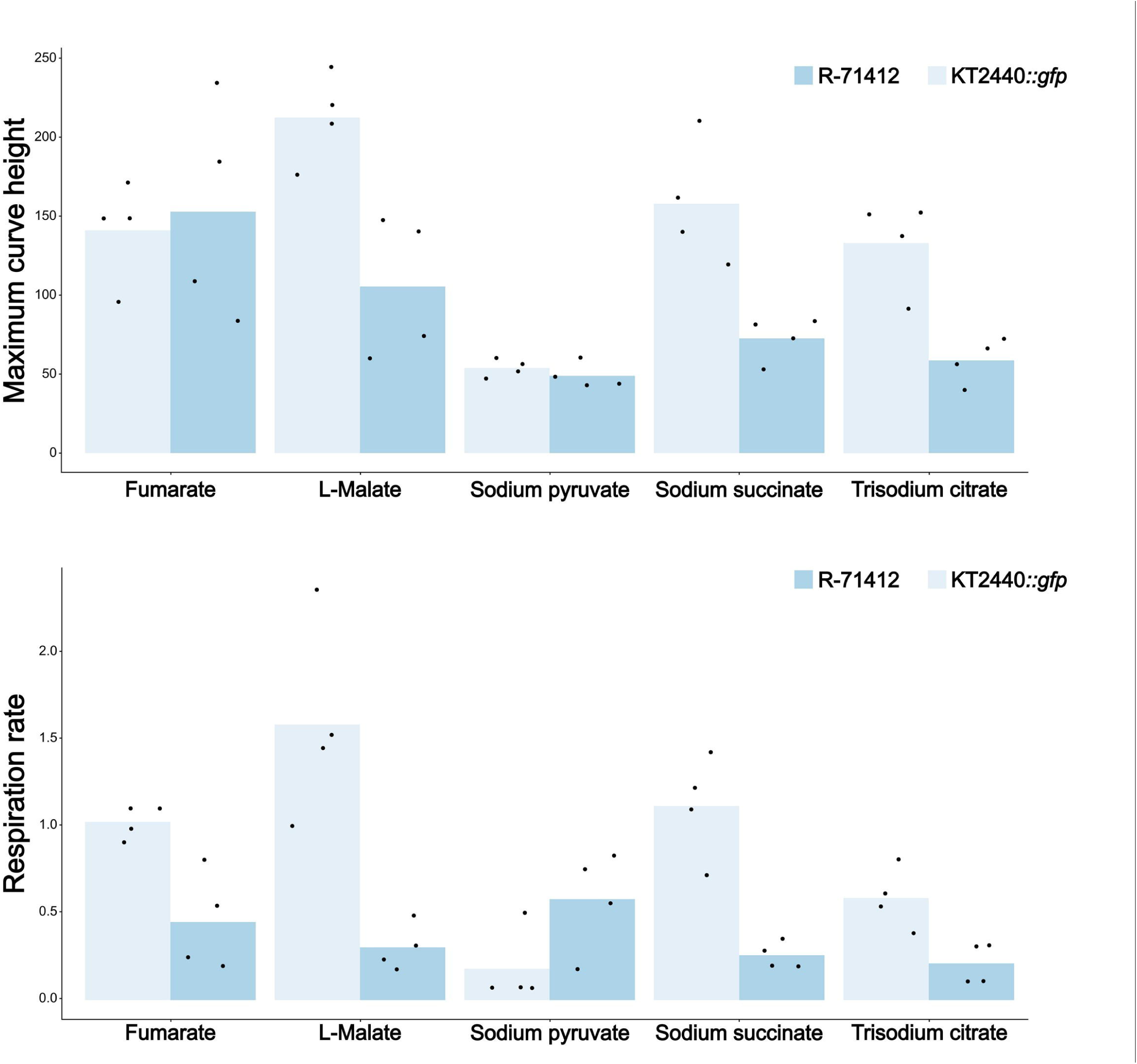
Growth characteristics of *P. putida* over O*. dioscoreae* on various carbon sources. Growth of *O. dioscoreae* R-71412 and *P. putida* KT2440*::gfp* was monitored on AB minimal medium supplemented with fumarate, L-malate, sodium pyruvate, sodium succinate and trisodium citrate during growth for 48 hours. The respiration rate µ and the Maximum curve height A were computed from curves modeled with R package Growthcurver.

To confirm that both strains compete for the same resources, we monitored growth of *O. dioscoreae* R-71417 and *P. putida* KT2440::*gfp* in co-cultures. The final cell density of *O. dioscoreae* R-71417 (mCherry-tagged) in LB medium (as measured via relative RFP-specific fluorescence) was drastically decreased in co-cultures with *P. putida* KT2440::*gfp* (Figure 3). Maximal growth of *P. putida* (as measured by relative GFP-specific fluorescence) was also impacted by the presence of *O. dioscoreae* but to a lesser extent. Moreover, when *O. dioscoreae* R-71417 was inoculated before *P. putida*, the final cell density of *P. putida* KT2440::*gfp* decreased with the time of pre-conditioning by *O. dioscoreae*. These results indicate that *P. putida* and *O. dioscoreae* do compete for resources in liquid cultures, and that *P. putida* KT2440::*gfp* tends to consistently outgrow *O. dioscoreae*.

**Figure 3.**
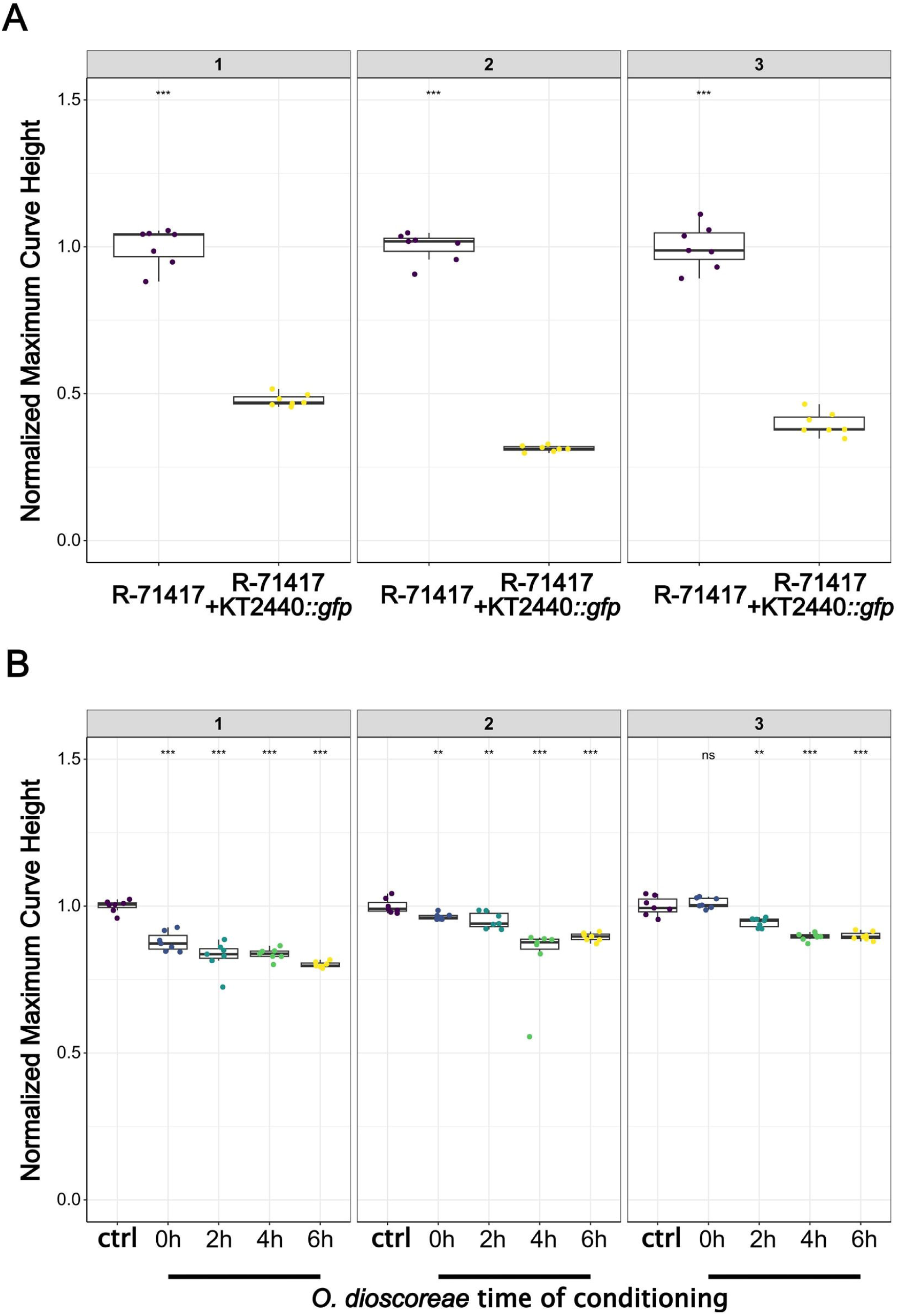
*In vitro* competition between *P. putida* and *O. dioscoreae* . The competition for resources was tested by co-culturing *P. putida* KT2440*::gfp* and mCherrry –tagged *O. dioscoreae* R-71417. LB medium was inoculated with mCherrry–tagged *O. dioscoreae* R-71417 before (2, 4 or 6h) or simultaneously with *P. putida* KT2440*::gfp*. Controls (Ctrl) correspond to single cultures of *P. putida*. (A) GFP and (B) mCherry fluorescence intensity were monitored for 24 hours. The maximum height of the growth curve was calculated using R package Growthcurver, and for each individual replicate the maximum height was normalized based on the mean maximum height of the single cultures. Results of independent experiments are shown as separate panels. Statistical significance calculated with a Wilcoxon test (significance levels: ns p-value >0.05; * p-value <=0.05; ** p-value <=0.01; *** p-value <=0.001; **** p-value <=0.0001).

### Effective colonization of *D. sansibarensis* leaf glands by *P. putida*

To test whether *P. putida* KT2440 could establish stable infections in *D. sansibarensis*, we infected aposymbiotic plantlets with *P. putida* KT2440::*gfp*. We detected *P. putida* KT2440::*gfp* within the glands of all plants infected with the strain, but in none of the mock-inoculated controls. Leaf glands of plants inoculated with strain KT2440::*gfp* contained an average of 10^9^ cfu/g of fresh tissue (Figure 5A), while lamina of surface-sterilized leaves did not contain detectable bacteria. This is similar to the colonization levels of wild-type *O. dioscoreae* in leaf glands of *D. sansibarensis* (Figure 1). Epifluorescence imaging of leaf forerunner tip sections further confirmed that cells of *P. putida* KT2440::*gfp* occupied the lumen of the leaf glands, similar to *O. dioscoreae* in wild-type plants (Figure 4).

**Figure 4.**
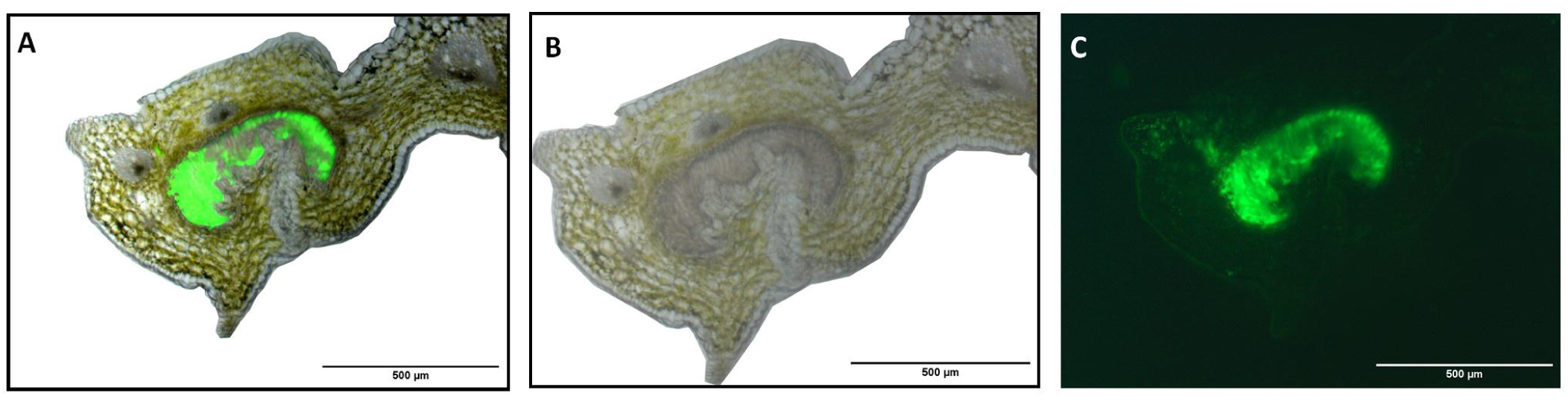
Localization of *P. putida* KT2440 to *D. sansibarensis* leaf glands. A. Cross-section of *D. sansibarensis* acumen colonized by *P. putida* KT2440*::gfp* after artificially inoculation. Merged image of light microscopy (B) and epifluorescence (C) observations.

**Figure 5.**
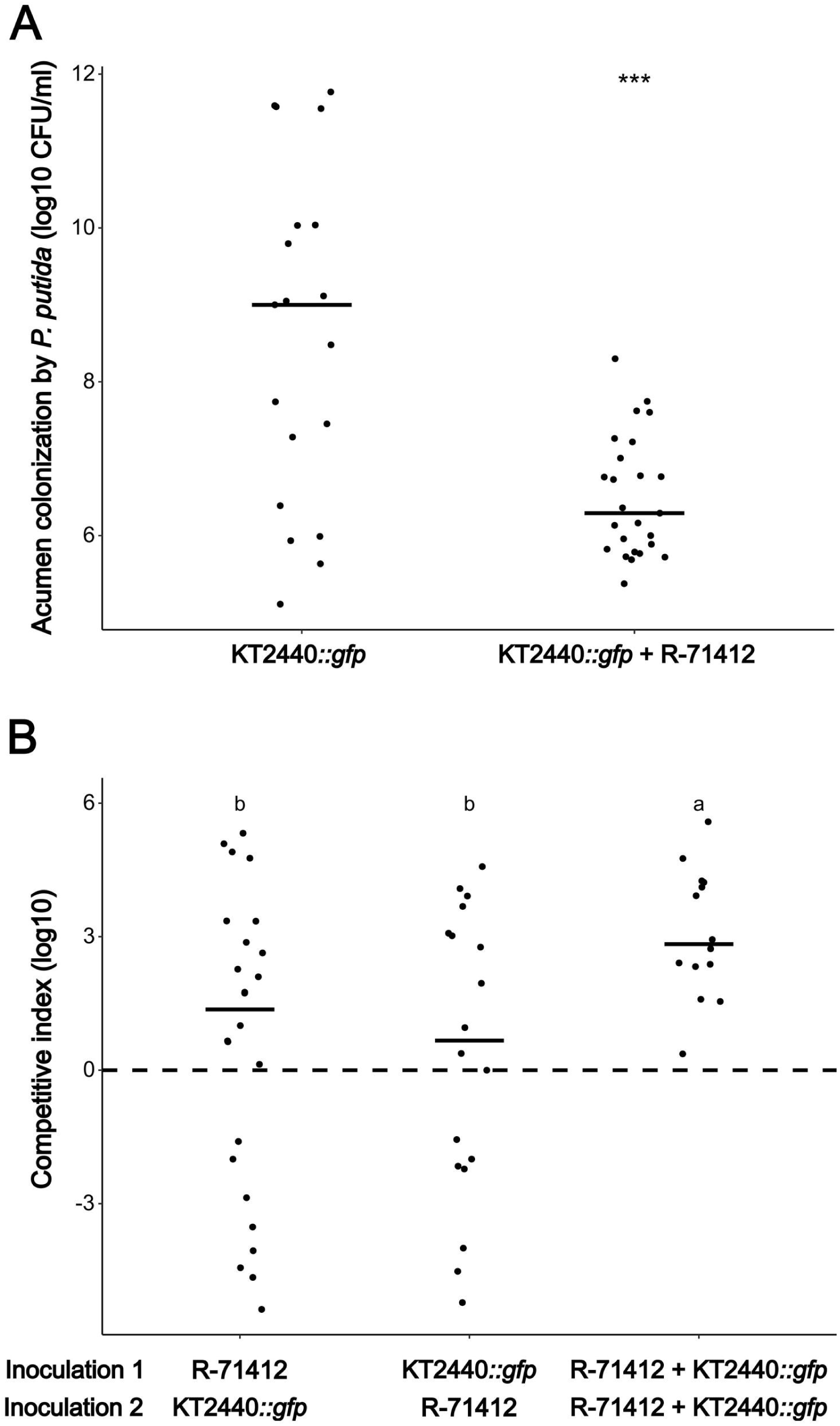
*In planta* competition between *O. dioscoreae* and *P. putida*. (A) Quantification of *P. putida* KT2440*::gfp* in leaf glands of *D. sansibarensis* after single inoculation or co-inoculation with *O. dioscoreae* R-71412 (1:1 ratio). Statistical significance calculated with Student’s t-test (significance levels: ns p-value >0.05; * p-value <=0.05; ** p-value <=0.01; *** p-value <=0.001; **** p-value <=0.0001). (B) Competitive index of *O. dioscoreae* R-71417 in successive co-inoculations with *P. putida* KT2440. Aposymbiotic plants were inoculated successively with either *O. dioscoreae* R-71412 or *P. putida* KT2440*::gfp*, or with the same strain twice. Competitive index was calculated as the log10 of the acumen colonization (cfu per g) ratio R-71412:KT2440. Different letters indicate statistically significant differences between groups according to ANOVA with Tukey post-hoc test (significance threshold *p* = 0.05).

### Competitive advantage of *O. dioscoreae* in the leaf gland

Since colonization of leaf glands did not seem to be specific to *O. dioscoreae*, we wondered whether better niche adaptation of *O. dioscoreae*, or competitiveness, might account for specificity in nature. To test this, we co-inoculated aposymbiotic plants with cell suspensions of *P. putida* KT2440::*gfp* alone or mixed with *O. dioscoreae*. The presence of *O. dioscoreae* drastically reduced the average titer of *P. putida* KT2440::*gfp* in the leaf gland by a factor of 5x10^3^ (from 9.15x10^10^ to 1.67x10^7^ cfu/g) (Figure 5A).

Although co-culture experiments in liquid media may allow the detection of nutrient-based competition, they fail to account for several factors known to affect the outcome of competition. Niche conditioning by *O. dioscoreae*, e.g. by depleting nutrients, saturating favored spatial niches, secreting antimicrobial compounds or inducing specific plant defenses, might explain the lower rates of growth of commensal strains in the leaf gland in presence of *O. dioscoreae*. Manipulating the order of inoculation between the strains would thus be expected to exacerbate the differences.

To investigate the role of the competition in the ability of the strains to colonize the plant, we inoculated the two strains in varying order: *O. dioscoreae* first, *P. putida* KT2440::*gfp* first or both simultaneously. We did not observe any significant effect of the order of arrival on the competitive index of either strain in the leaf gland, indicating that niche conditioning or priority effects do not account for the majority of the competitive advantage of *O. dioscoreae* in the leaf gland (Figure 5B). However, co-inoculations always resulted in a significant competitive advantage of *O. dioscoreae*.

### Competitive advantage of *O. dioscoreae* is mediated by T6SSs

Because the genome of *O. dioscoreae* encodes two Type VI secretion systems, and that both are overexpressed *in planta* compared to *O. dioscoreae* grown on minimal growth medium [22], we reasoned that antimicrobial effectors secreted by T6SS may give a competitive advantage to *O. dioscoreae in planta*. We tested contact-dependent growth inhibition of *P. putida* KT2440::*gfp* by *O. dioscoreae* in a microtiter plate assay. We observed a significant decrease of GFP-specific fluorescence in the presence of *O. dioscoreae* R-71412. This effect was reduced in the presence of strains harboring mutations in either or both of the T6SS *clpV* genes (Figure 6). Complementation of the single and double mutant with copies of *clpV1* or *clpV2 in trans* restored the wild-type phenotype.

**Figure 6.**
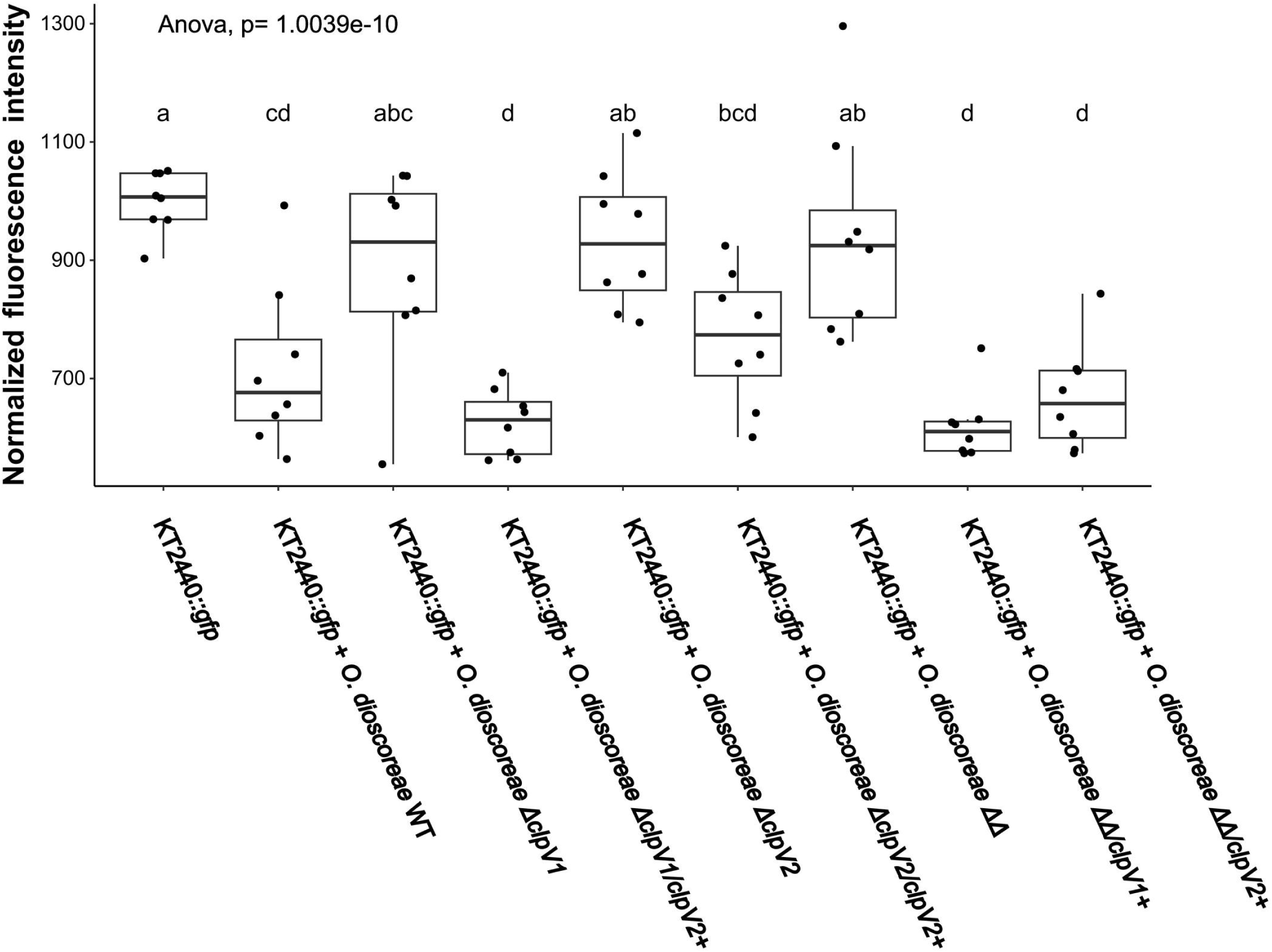
Contact-dependent competition assays between *O. dioscoreae* against *P. putida*. Normalized fluorescence intensity of *P. putida* KT2440*::gfp* after 4 hours of growth alone or in contact with *O. dioscoreae* strains. *O. dioscoreae* T6SS mutants *ΔclpV1*, *ΔclpV2* and *ΔclpV1ΔclpV2* and complemented strains *ΔclpV1/clpV1+*, *ΔclpV2/clpV2+*, *ΔclpV1ΔclpV2/clpV1+* , and *ΔclpV1ΔclpV2/clpV2+* were tested. Different letters indicate statistically significant differences between groups according to ANOVA with Tukey post-hoc test (significance threshold *p* = 0.05). Results shown here are from one of three independent experiments showing similar results.

To confirm that contact-dependent killing is responsible for the reduction in GFP-specific fluorescence in the microtiter plate assay, we counted the number of cfu by dilution plating of both *P. putida* KT2440::*gfp* and *O. dioscoreae* strains after direct contact. We observed a stark decrease in the number of *P. putida* KT2440::*gfp* colonies after 4 hours of contact with *O. dioscoreae* R-71412, but not with the *ΔclpV1ΔclpV2* mutant (Figure S3).

Finally, we designed a co-inoculation experiment to test the contribution of T6SS to the competitive advantage of *O. dioscoreae in planta*. The presence of the wild-type strain of *O. dioscoreae* negatively impacted the colonization of *P. putida* KT2440::*gfp* by several orders of magnitude as observed previously. In contrast, this negative effect was much alleviated when co-inoculating plants with the *O. dioscoreae ΔclpV1ΔclpV2* strain (Figure 7).

**Figure 7.**
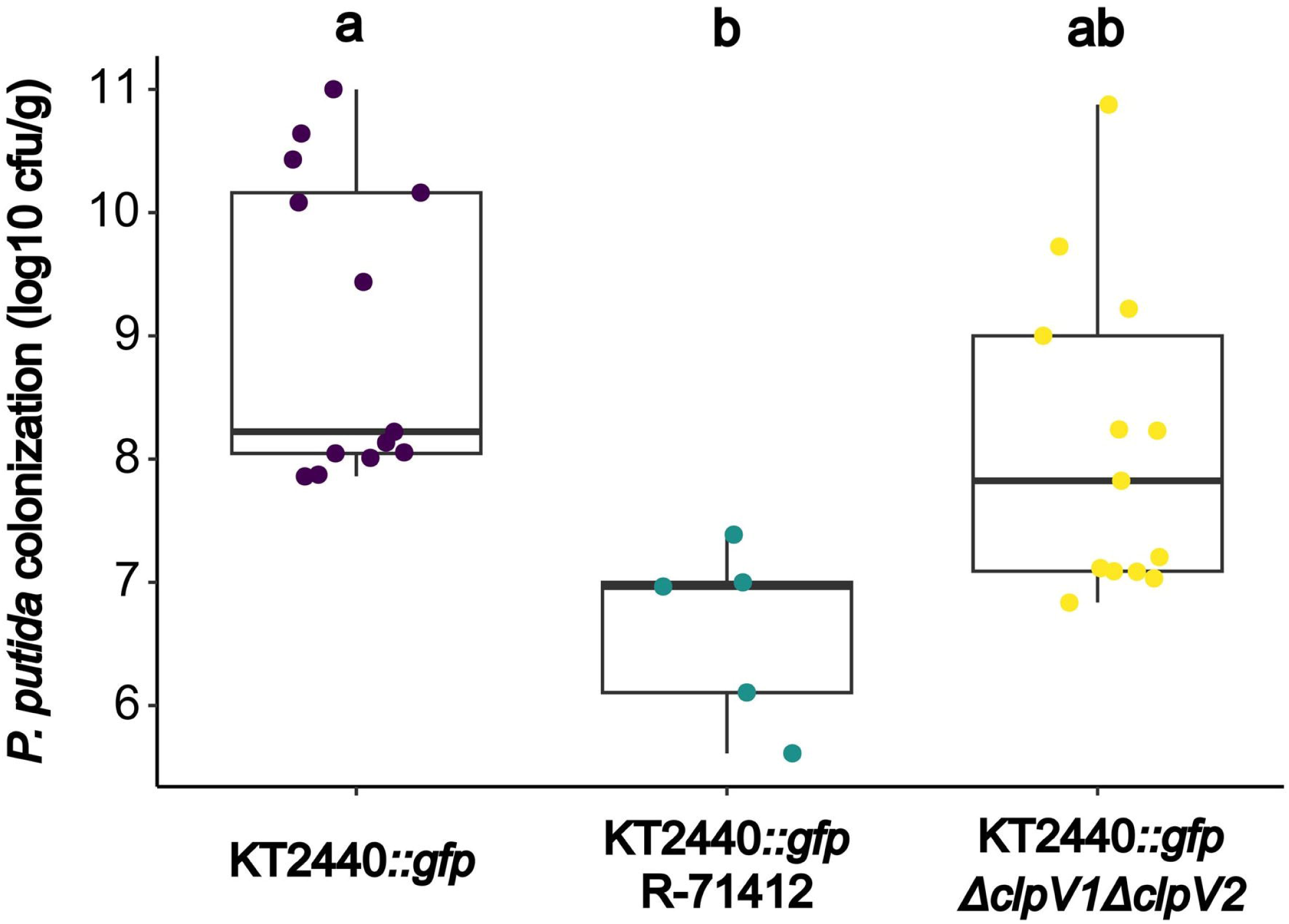
*In planta* competition between P. putida and strains of *O. dioscoreae* affected in Type VI secretion. Aposymbiotic plants were inoculated with *P. putida* KT2440*::gfp* alone or in 1:1 ratio with *O. dioscoreae* strain R-71412, or strain *ΔclpV1ΔclpV2*. Freshly grown acumens were weighed and macerated. Serial dilutions were plated on selective media to quantify bacterial load. Different letters indicate statistically significant differences between groups according to ANOVA with Tukey post-hoc test (significance threshold *p* = 0.05). Results shown here are from one of two independent experiments showing similar results.

### Evidence of species-selective transmission

We wondered whether *P. putida* could also be transmitted to the next generation of plants. To test this, we inoculated plants with *P. putida* KT2440::*gfp*. We collected the bulbils produced by the plants at the end of the growing season and let them germinate after a rest period. The plants produced on average 5.9 bulbils per plant, with a bulbil germination rate of 9.1% (Table 3). None of the daughter plants contained detectable *P. putida* in the leaf glands. As a control, we also inoculated plants with a 1:1 cell mixture of *P. putida* KT2440::*gfp* and *O. dioscoreae* R-71412. These produced 4.4 bulbils by plant on average, for a bulbil germination rate of 15.7%. Of the 109 daughter leaf acumens tested only 1 contained both *P. putida* KT2440::*gfp* and *O. dioscoreae* (0.9% of the leaf glands), while we detected *O. dioscoreae* R-71412 in 69 samples (63.3%).

## Discussion

Heritable leaf symbioses are highly specific in nature: one plant species associated with only one bacterial species. At least in the case of the *Dioscorea sansibarensis* leaf symbiosis, this specificity is not due to an obligate interaction: aposymbiotic plants develop normally in a controlled environment, and bacteria can grow separately as well [17]. Several recent studies have shown that free-living bacteria could effectively replace vertically-transmitted insect symbionts, under some conditions. For example, the obligate symbiont *Sodalis pierantonius* of the grain weevil *Sitophilus zeamais* could be replaced by a free-living strain of *Sodalis praecaptivus* engineered to secrete aromatic amino acids [51]. Similarly, a free-living *E. coli* strain was successfully evolved in the laboratory and provided an effective replacement for the vertically-transmitted *Pantoea* symbiont of *Plautia stali* stinkbugs upon acquisition of a mutation in the global regulators controlling carbon utilization pathways [52]. In both of these examples, stable symbiont replacement occurred with closely-related strains (same genus or family), and after some genetic modification. Here, we show that a vertically-transmitted plant symbiosis is surprisingly promiscuous, with taxonomically diverse strains capable of stable colonization. This is in stark contrast to the 100% specificity for *O. dioscoreae* observed in field samples [18, 22].

Using metabolomics data from *D. sansibarensis* leaf glands and metabolic profiling of *O. dioscoreae*, we established the conditions likely encountered by the bacteria inside the plant. Conditions in the *D. sansibarensis* leaf glands are micro-oxic (De Meyer et al 2019), and *O. dioscoreae* grows best at neutral to slightly acidic pH. This could indicate that tolerance to acid and oxidative stress is a requirement for life inside the plant. Additionally, *O. dioscoreae* putative high-affinity iron-acquisition systems are overexpressed *in planta* compared to axenic cultures [22]. This is perhaps indicative of depletion of iron in the leaf gland, which may be associated with plant immunity [53]. Furthermore, genes involved in oxidative stress responses (e.g. putative catalase and peroxidases) are overexpressed by the bacteria *in planta*, and functions linked to resistance to oxidative stress are highly conserved in the core genomes of leaf symbionts [22, 54]. TCA cycle substrates could be commonly use as major carbon sources by leaf endosymbionts. Taken together, these data indicate that the conditions in the leaf glands may contribute to selecting bacterial specialists adapted for utilization of acids and tolerance to oxidative stress, but may not be otherwise particularly stringent.

Indeed, *P. putida* KT2440 presents an overlapping metabolic profile to *O. dioscoreae*, and is able to colonize the leaf glands of *D. sansibarensis* to high titer. *P. putida* KT2440 derives from a soil isolate and is unlikely to be pre-adapted to an endophytic lifestyle. This suggests that once the plant is deprived of its symbiont the leaf gland became open to colonization by other bacterial strains.

In *D. sansibarensis* glands, bacteria are present to high titer up to 1x10^10^ cfu/g of tissue, at least an order of magnitude higher than what is achievable in liquid cultures. High cell-density and proximity allows for contact-dependent competition. Experiments on solid media indeed showed that *O. dioscoreae* kills *P. putida* KT2440::*gfp* cells in a contact-dependent manner, and that this antagonistic activity is mediated by 2 T6SS gene clusters encoded in the genome of *O. dioscoreae.* Both T6SS-1 and T6SS-2 of *O. dioscoreae* LMG29303^T^ are overexpressed *in planta* and are conserved in all *O. dioscoreae* genomes, although the repertoire of predicted effectors vary between strains [18, 22]. The antimicrobial activity of T6SS and associated effectors is well-described, and contrasting T6SS effector sets may reflect varying pathogen or competitor pressures in natural populations of *O. dioscoreae* (reviewed in [32, 55]). Importantly, we show that the T6SSs of *O. dioscoreae* contribute to the advantage *in planta* against *P. putida*, but do not affect the ability to colonize the leaf gland in single inoculations (Figure S4). Aside from a role in protecting the symbiotic niche, it is unclear whether *O. dioscoreae* also provides protection against bacterial or fungal pathogens, as has been demonstrated in other systems [56, 57].

Interestingly, complete T6SS were only detected in the genomes of leaf symbionts of *Fadogia homblei* and *Vangueria pygmaea*, but not in other leaf symbionts of the *Caballeronia* clade [11]. How specificity is maintained in leaf symbioses in the absence of a T6SS remains unknown, but other secreted toxins might play a role.

Alone, antagonistic microbe-microbe interactions are unlikely to fully explain the absolute specificity between *D. sansibarensis* and *O. dioscoreae* observed in the wild. The absence of a phenotype in aposymbiotic plants suggests that symbiotic functions may be important for fitness in response to environmental or herbivory pressures. We are currently assessing the phenotypes of plants under an array of conditions. If the symbiosis is indeed essential for survival under natural conditions, vertical transmission and partner-fidelity feedback may also contribute to specificity by culling plants with ineffective symbionts from the population [58, 59].

Finally, however promiscuous *D. sansibarensis* is in artificial infections, transmission of symbionts to bulbils appears more stringent. Transmission appears to be imperfect in our experiments, with bacteria detected in only 62% of bulbils. This indicates that barriers to transmission exist. These barriers appear somewhat selective, with plants inoculated with *P. putida* alone unable to pass on the bacterium to their offspring. Only when we co- inoculated plants with *O. dioscoreae* did we detect *P. putida* in the offspring of plants, albeit at anecdotal frequencies. It is unclear what the mechanisms enforcing specificity of transmission might be, but we speculate that the ability to withstand a quiescent phase of several months before bulbils germinate might be an important factor in transmission success.

This study clearly emphasizes the complexity of specificity regulation in the *Dioscorea- Orrella* symbiosis. The absence of a strong phenotype of aposymbiotic plants, the remarkable promiscuity of the plant for bacteria in artificial infections, and the fact that specificity seems enforced (at least partially) by microbe-microbe interactions suggest that the barriers to evolving vertically-transmitted plant symbioses may be surprisingly low. This remarkable plasticity may explain why leaf symbiosis seems to have evolved independently several times in distinct plant lineages. What adaptations underly the evolution of vertical transmission in these plant-bacteria associations remain to be discovered.

## ACKNOWLEDGMENTS

LN and AC acknowledge support by the French Laboratory of Excellence project "TULIP" (ANR-10-LABX-41; ANR-11-IDEX-0002-02). This study is set within the framework of the ‘École Universitaire de Recherche (EUR)’ TULIP-GS (ANR-18-EURE-0019). AC also wishes to acknowledge funding from the French National Research Agency under grant agreements ANR-22-CE92-0042 and ANR-23-CE11-0015, and the Région Occitanie through the “RAMSY” project. Financial support was received from the French National Infrastructure for Metabolomics and Fluxomics, Grant MetaboHUB-ANR-11-INBS-0010. We wish to thank Nicolas Krink and Pablo Nikel from the Systems Environmental Microbiology Group of the Novo Nordisk Foundation Center for Biosustainability at DTU (Denmark Technical University) for the kind gift of *P. putida* strains.

## SUPPLEMENTARY FIGURES AND TABLES

FIG S1, Symbiotic and non-symbiotic bacteria localize to Dioscorea sansibarensis leaf glands.

FIG S2, Growth of O. dioscoreae R-71412 at various pH.

FIG S3, Contact-dependent competition assays of O. dioscoreae against P. putida KT2440.

FIG S4, D. sansibarensis acumen colonization by O. dioscoreae strains.

TABLE S1, Strains and Plasmids.

TABLE S2, Oligonucleotides.

TABLE S3. Metabolomic profile of symbiotic and aposymbiotic D. sansibarensis leaf glands.

TABLE S4. Annotated features detected in extracts of symbiotic and aposymbiotic leaf acumens of D. sansibarensis measured by UHPLC-HRMS.

