## Supplementary figures for "Artificial symbiont replacement in a vertically-transmitted plant-bacterium association provides insights into the basis for specificity"

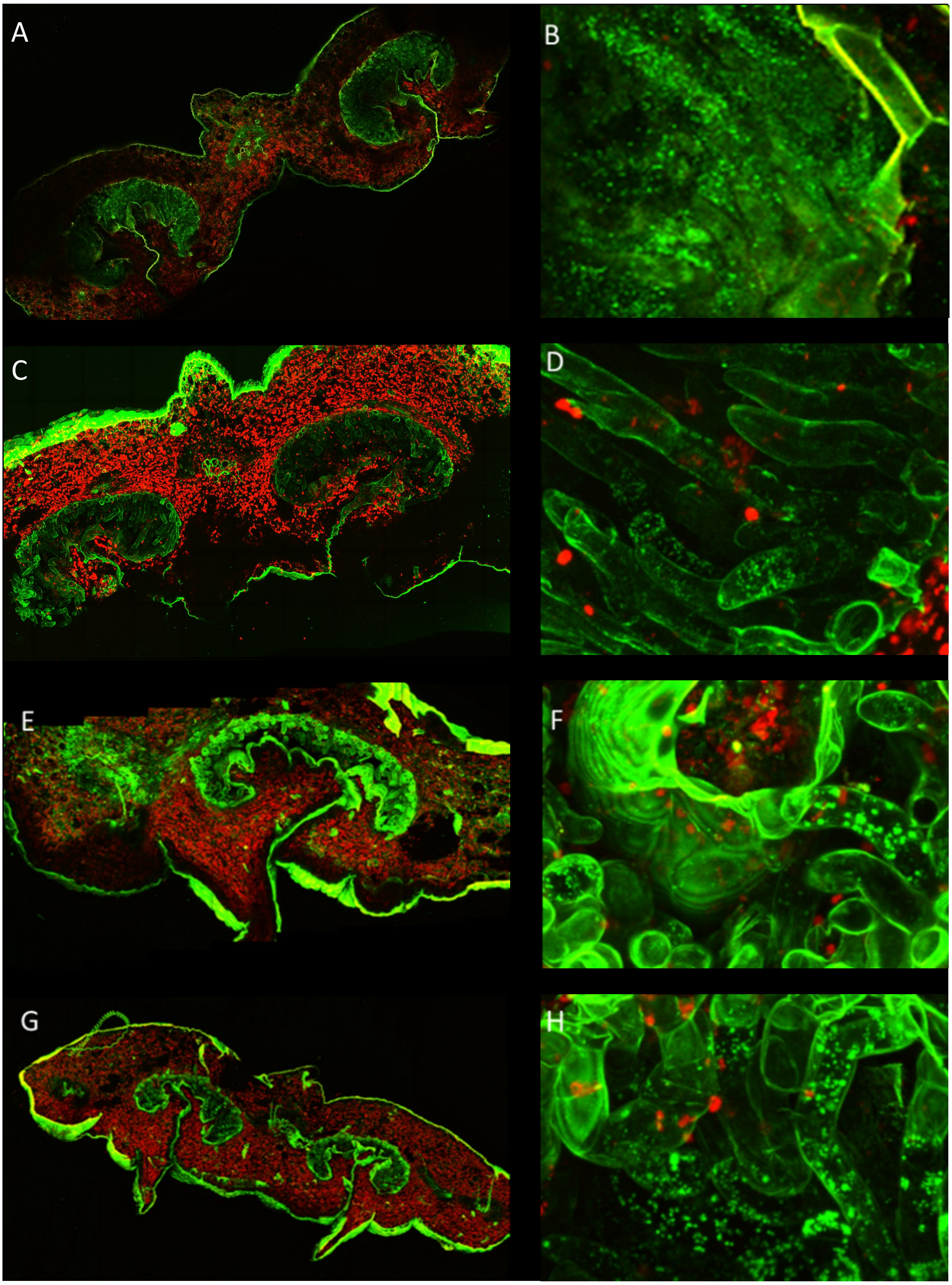


**Figure S1 Symbiotic and non-symbiotic bacteria localize to *Dioscorea sansibarensis* leaf glands.** Confocal images of cross sections of *D. sansibarensis* leaf acumens colonized by different bacterial species after artificial infection. *D. sansibarensis* leaf glands colonized by (A) *O. dioscoreae*, (C) *Rhizobium sp.*, (E) *Stenotrophomonas sp.* and (G) *Sphingomonas sp.*. Details of trichomes surrounded by mucus and green foci corresponding to stainec bacteria (B) *O. dioscoreae*, (D) *Rhizobium sp*, (F) *Stenotrophomonas sp.* and (H) *Sphingomonas sp.*. Bacteria were stained with Syto9 and red autofluorescence was used to visualize plant cells.

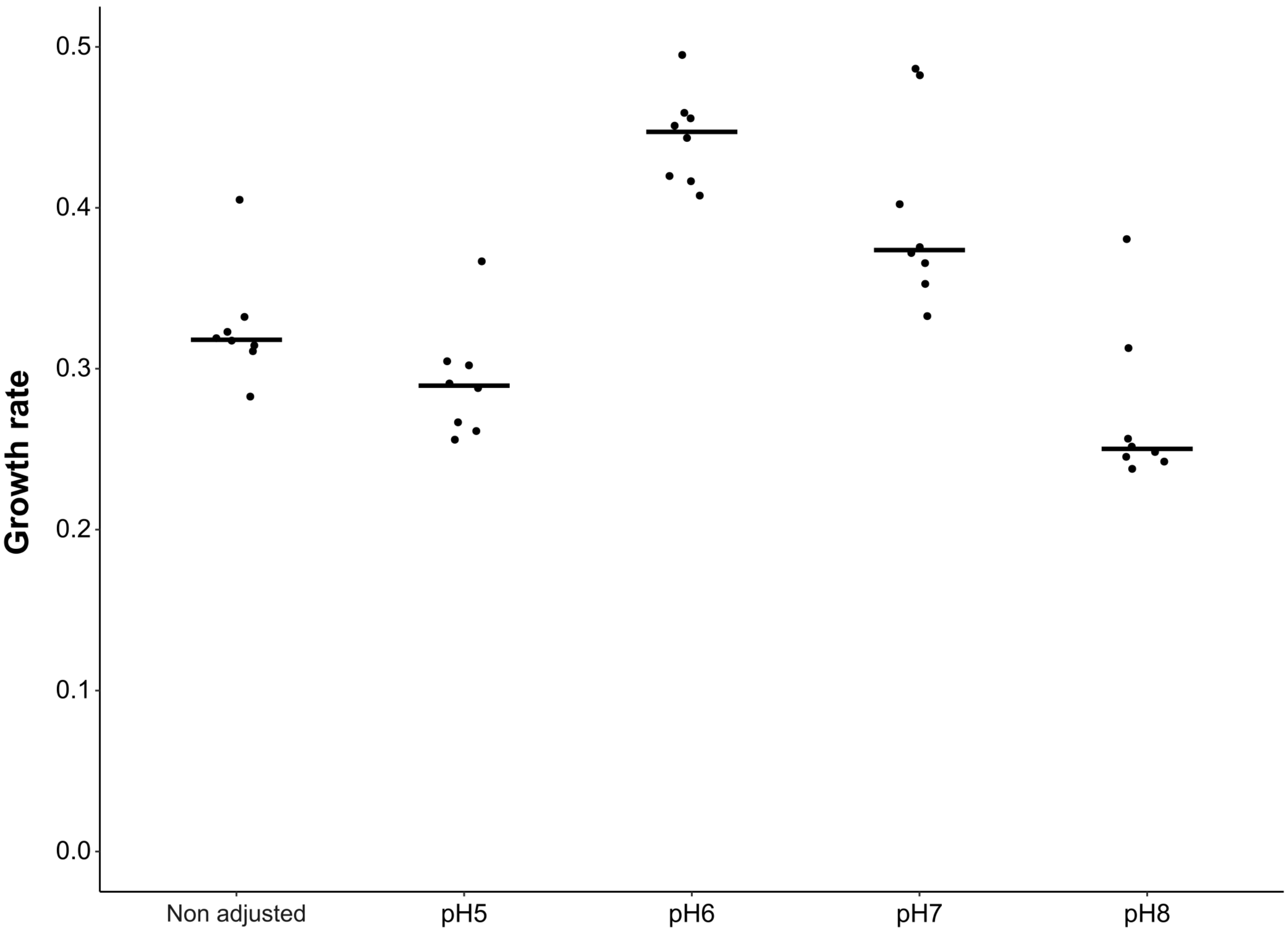


**Figure S2. Growth of *O. dioscoreae* grows at various pH.** *O. dioscoreae* R-71417 growth was monitored for 48 hours in LB medium supplemented with 10mM trisodium citrate from pH5 to pH8. The pH was adjusted to and buffered with MES, MOPS and EPPS buffer; non adjusted medium was used as a control. Growth rate was calculated using R package Growthcurver (Sprouffske et Wagner 2016).

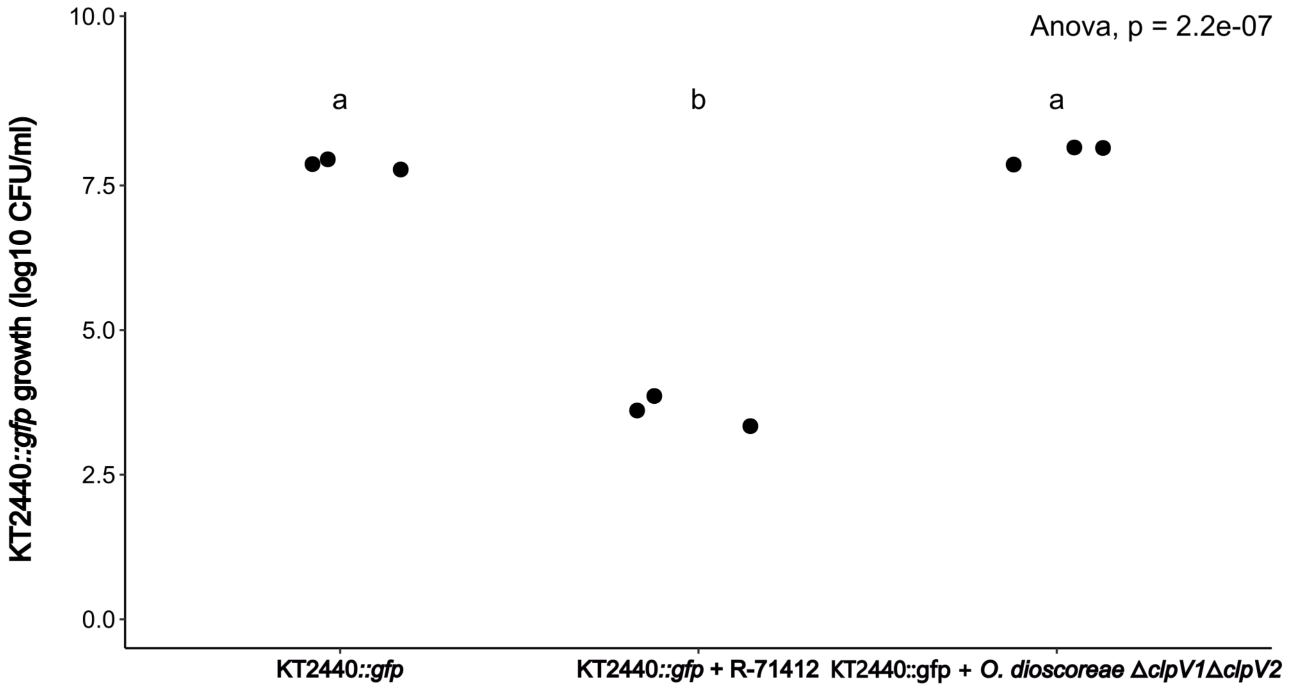


**Figure S3. Contact-dependent competition assays of *O. dioscoreae* against *P. putida* KT2440::*gfp*.** *O. dioscoreae* T6SS-mediated competition against *P. putida* KT2440::gfp. After 4 hours of growth on TSA medium alone or mixed

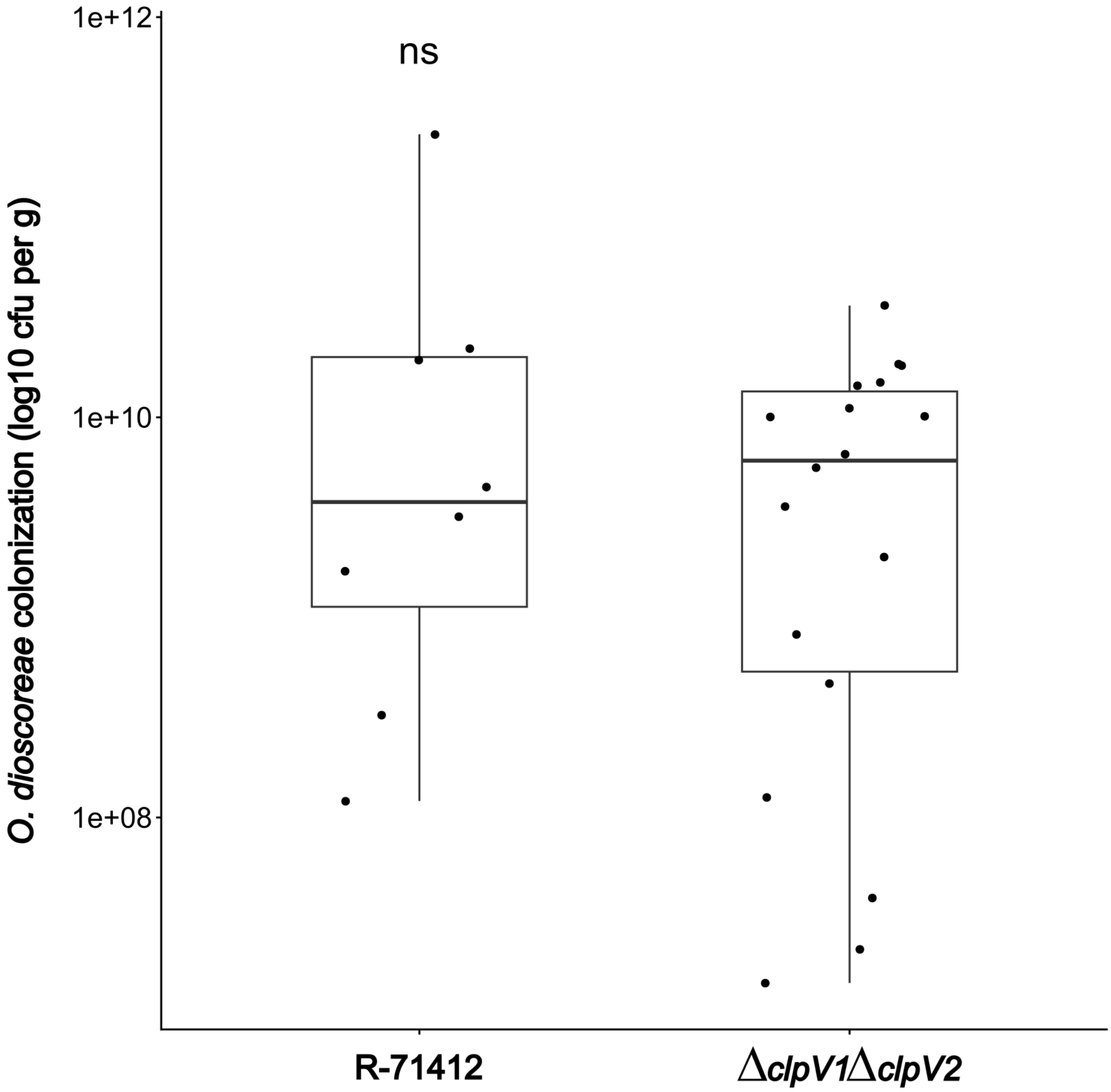


**Figure S4.** ***D. sansibarensis*** **acumen colonization by** ***O. dioscoreae*** **strains.** Aposymbiotic *D. sansibarensis* were inoculated with *O. dioscoreae* strains R-71412, the T6SS-mutant *ΔclpV1ΔclpV2*. Newly grown acumens were weighed and macerated. Serial dilutions were plated on selective media to quantify bacterial load. Statistical significance calculated with Student's t-test (significance levels: ns p-value >0.05; * p-value <=0.05; ** p-value <=0.01; *** p-value <=0.001; **** p-value <=0.0001).
