## Supplementary material for "Artificial symbiont replacement in a vertically-transmitted plant-bacterium association provides insights into the basis for specificity": Table S1

**Table S1 Strains and Plasmids**

| Species | Strain or plasmid | Description | Growth conditions | Reference or source |
| --- | --- | --- | --- | --- |
|  | Strain |  |  |  |
| *Orrella dioscoreae* | R-71412 | Spontaneous nalidixic acid resistant strain derived from type strain LMG 29303^T^. | TSA + Nalidixic acid 30 µg/ml, aerobic, 28°C | (De Meyer et al. 2019) |
| *Orrella dioscoreae* | R-71417 | mCherry-tagged derivative of R-71412 | TSA + Nalidixic acid 30 µg/ml + Gentamicin 20 µg/ml, aerobic, 28°C | (Acar et al. 2022) |
| *Escherichia coli* | Top10 |  | LB, 37°C | Thermo Fisher Scientific |
| *Orrella dioscoreae* | ∆3997 | Deletion mutant in *clpV1* gene of , derivative of strain R-71412 | TSA + Nalidixic acid 30 µg/ml, aerobic, 28°C | This study |
| *Orrella dioscoreae* | ∆0808 | deletion mutant in *clpV2* gene, derivative of R-71412 | TSA + Nalidixic acid 30 µg/ml, aerobic, 28°C | This study |
| *Orrella dioscoreae* | ∆3997∆0808 | deletion mutant in *clpV1* and *clpV2* genes, derivative of R-71412 | TSA + Nalidixic acid 30 µg/ml, aerobic, 28°C | This study |
| *Orrella dioscoreae* | ∆3997/*clpV1+* | pSEVA2313_R3997 | TSA + Nalidixic acid 30 µg/ml + Kanamycin 50 µg/ml, aerobic, 28°C | This study |
| *Orrella dioscoreae* | ∆0808/*clpV2+* | pSEVA2313_R0808 | TSA + Nalidixic acid 30 µg/ml + Kanamycin 50 µg/ml, aerobic, 28°C | This study |
| *Orrella dioscoreae* | ∆3997∆0808/*clpV1+* | pSEVA2313_R3997 | TSA + Nalidixic acid 30 µg/ml + Kanamycin 50 µg/ml, aerobic, 28°C | This study |
| *Orrella dioscoreae* | ∆3997∆0808/*clpV2+* | pSEVA2313_R0808 | TSA + Nalidixic acid 30 µg/ml + Kanamycin 50µg/ml, aerobic, 28°C | This study |
| *Pseudomonas putida* | KT2440::*gfp* | P_14g_(BCD2)->msfGFP (monomeric superfolder GFP under control of a P14g promoter and a biscistronic design), chromosomally integrated into a landing pad between genes PP_0013 and PP_5421 | LB, 28°C | Gift from Nicolas Krink (Nikel lab DTU) |
|  | Plasmid |  |  |  |
| *Escherichia coli* | pQURE6 | Top10 | LB + Gentamicin 20 µg/ml, 37°C | (Volke, Wirth, et Nikel 2021) |
| *Escherichia coli* | pSEVA2313 | CC118 | LB + Kanamycin 25 µg/ml, 37°C | (Silva-Rocha et al. 2013) |
| *Escherichia coli* | pSNW2 | DB3.1 lambda pir |  | (Volke, Wirth, et Nikel 2021) |
| *Escherichia coli* | pUX-BF13 | SM10 lambda pir | LB + Ampicillin 100 µg/ml, 37°C | (Choi et Schweizer 2005) |
| *Escherichia coli* | pSNW2_mut_R0808 | DB3.1 lambda pir |  | This study |
| *Escherichia coli* | pSNW2_mut_R3997 | DB3.1 lambda pir |  | This study |
| *Escherichia coli* | pSEVA2313_R0808 | Top10 | LB + Kanamycin 25µg/ml, 37°C | This study |
| *Escherichia coli* | pSEVA2313_R3997 | Top10 | LB + Kanamycin 25µg/ml, 37°C | This study |
