## Supplementary material for "Artificial symbiont replacement in a vertically-transmitted plant-bacterium association provides insights into the basis for specificity": Table S2

**Table S2 Oligonucleotides**

| Primers | Sequence | Description |
| --- | --- | --- |
| **27F** | AGAGTTTGATCCTGGCTCAG | 16S sequencing |
| **1492R** | TACGGYTACCTTGTTACGACTT |  |
| **R3997_L_fwd** | TGAATTCGAGCTCGGTACCCGCGGGCTCGAGCAGCTGGGC | Deletion of ODI_R3997 |
| **R3997_L_rev** | TCACAGAGCCCAGCGCCATGTGACACGGGAGCCACGCCAT |  |
| **R3997_R_fwd** | ATGGCGTGGCTCCCGTGTCACATGGCGCTGGGCTCTGTGA |  |
| **R3997_R_rev** | GTCGACTCTAGAGGATCCCCTGCGCATCCAGGAGTTCGTG |  |
| **R0808_L_fwd** | TGAATTCGAGCTCGGTACCCGCGCGTGCTGTCCGACTACT | Deletion of ODI_R0808 |
| **R0808_L_rev** | GCTGGTGCTCCCTCAGCTCACATGGAAGGTTGCGTCGGGG |  |
| **R0808_R_fwd** | CCCCGACGCAACCTTCCATGTGAGCTGAGGGAGCACCAGC |  |
| **R0808_R_rev** | GTCGACTCTAGAGGATCCCCCGGTCCTTCACTTCCCAGCG |  |
| **R3997_fwd_EcoRI** | GGGGAATTCTCACAGAGCCCAGCGCCATG | Trans complementation mutant of ODI_R3997 |
| **R3997_rev_HindIII** | GGGAAGCTTTCAGGCGAAGTCGAAGGTGA |  |
| **R0808_fwd_EcoRI** | GGGGAATTCGCCATCACCGAACCCCG | Trans complementation mutant of ODI_R0808 |
| **R0808_rev_HindIII** | GGGAAGCTTTCAGCTCACGCGGTAGCGGAAC |  |
